# Abundance of Nef and p-Tau217 in brains of individuals diagnosed with HIV-associated neurocognitive disorders correlate with disease severance

**DOI:** 10.1101/2021.07.20.453134

**Authors:** Tatiana Pushkarsky, Adam Ward, Dmitri Sviridov, Michael I Bukrinsky

## Abstract

HIV-associated neurological disorders (HAND) is a term used to describe a variety of neurological impairments observed in HIV-infected individuals. The pathogenic mechanisms of HAND and its connection to HIV infection remain unknown. The brain samples from HIV-infected individuals, both with and without HAND, were characterized by increased abundance of p-Tau217 peptide, which correlated with the abundance of flotillin 1, a marker of lipid rafts. HIV-1 Nef was detected in some, but not all, samples from HAND-affected individuals. Samples positive for Nef had lower abundance of cholesterol transporter ABCA1, higher abundance of flotillin 1 and p-Tau217, and were obtained from individuals with higher severity of HAND relative to Nef-negative samples. These results highlight the contribution of Nef and Nef-dependent effects on cholesterol metabolism and lipid rafts to the pathogenesis of HAND and support a connection between pathogenesis of HAND and Alzheimer’s disease.

## Introduction

HIV-associated neurological disorders (HAND) have clinical hallmarks of a neurodegenerative disease with progressive cognitive decline reflected initially by neuronal dendritic simplification and ultimately by the neuronal loss [1]. With the introduction of combination anti-retroviral therapy (cART), prevalence of the severe form of HAND, HIV-associated dementia (HAD), has diminished, but that of milder forms, mild neurocognitive disorder (MND) and especially asymptomatic neurocognitive impairment (ANI), have increased [2], suggesting that cART slows progression, but does not prevent initiation of the disease. The reasons for this persistence are not fully understood and several possible explanations have been suggested (reviewed in [3, 4]). One of the mechanisms gaining support in the recent studies links HAND pathogenesis to beta amyloid, suggesting a connection between HAND and Alzheimer’s disease (reviewed in [5]). HAND has many similarities to Alzheimer’s disease (AD), such as neuroinflammation, similar transcriptional signatures, and increased abundance and changed localization of intracellular beta amyloid (Aβ), a critical component in AD pathogenesis [5–8]. While the possibility that HIV infection “speeds up” the underlying AD is an attractive hypothesis, the mechanisms of this connection remain unclear. Described below are results of our previous studies of the effects of Nef on cellular cholesterol metabolism and lipid rafts, which provide a plausible mechanism.

Expression of Nef in HIV-infected cells induces degradation of the key cellular cholesterol transporter ABCA1, causing suppression of cholesterol efflux and increasing abundance of the lipid rafts [9, 10]. Most importantly, the same activity was demonstrated for the Nef-carrying extracellular vesicles [11, 12], which continue to be released into circulation and the brain from HIV-infected cells even after suppression of HIV replication by cART [7, 13, 14]. Amyloidogenic proteins concentrate in the lipid rafts [15], and most amyloid proteins involved in pathogenesis of neurodegeneration are raft proteins [16–18]. High local concentration of amyloidogenic proteins is a prerequisite for the effective nucleation and cascading progression of their misfolding, a key pathogenic element of neurodegeneration. Therefore, upregulation of the lipid rafts by Nef or Nef-containing extracellular vesicles and accumulation of amyloidogenic proteins in the rafts may accelerate the progress of their misfolding and promote its spread through the brain. A similar mechanism has been shown for prions, which also promote lipid raft formation and depend on lipid rafts for progression of prion misfolding [19].

In this study, we demonstrated associations between the presence of Nef with reduced levels of ABCA1, increased abundance of flotillin 1, disease severity, and the increased abundance of p-Tau217 - which is a characteristic marker of the Alzheimer’s disease [20–22]. Our findings support the proposed mechanism of HAND whereby Nef-mediated suppression of ABCA1 increases the abundance of lipid rafts, which in turn enables the progression of tauopathy.

## Materials and Methods

### Brain samples

Fresh-frozen post-mortem samples from hippocampus or mid-temporal gyrus of HIV-infected individuals with or without HAND diagnosis were obtained from National NeuroAIDS Tissue Consortium (NNTC). Brain samples from uninfected individuals without neurodegenerative disease diagnosis were obtained from the National Institutes of Health NeuroBioBank. All samples were deidentified, and provided clinical information is presented in Table S1. The studies abided by the Declaration of Helsinki principles.

### Western blot analysis

Samples were analyzed by automated Western immunoblotting using the JessTM Simple Western system (ProteinSimple, San Jose, CA). For analysis of ABCA1, a 66-440 kDa Jess separation module (SM-W008) was used, for Nef – a 2-40 kDa module (SM-W012), for p-Tau217 and Flotillin 1 – 12 - 230 kDa module (SM-W004). Three μL of brain lysates (1 μg/μL protein) was mixed with 1 μL of Fluorescent 5X Master mix (ProteinSimple) in the presence of fluorescent molecular weight markers and 400 mM dithiothreitol (ProteinSimple). This preparation was denatured at 95°C for 5 min for Nef, p-Tau217 and Flotillin 1 analysis. For ABCA1, the mix was incubated at 37°C for 15 min. Molecular weight ladder and proteins were separated in capillaries through a separation matrix at 375 volts. A ProteinSimple proprietary photoactivated capture chemistry was used to immobilize separated viral proteins on the capillaries. Capillaries with immobilized proteins were blocked with KPL Detection Block (5X) (SeraCare Life Sciences, Gaithersburg, MD, cat. #5920-0004) for 60 min, and then incubated with antibodies for 60 min. After a wash step, HRP-conjugated goat anti-rabbit (cat. #042-206), donkey anti-goat (cat. #043-522-2), or goat anti-mouse antibodies (cat. #042-205) from ProteinSimple were added for 30 min. The chemiluminescent revelation was established with peroxide/luminol-S (ProteinSimple). Digital image of chemiluminescence of the capillary was captured with Compass Simple Western software (version 5.1.0, Protein Simple) that calculated automatically heights (chemiluminescence intensity), area, and signal/noise ratio. Results were visualized as electropherograms representing peak of chemiluminescence intensity and as lane view from signal of chemiluminescence detected in the capillary. A total protein assay using Total Protein detection module DM-TP01 and Replex Module RP-001 was included in each run to quantitate loading.

### Antibodies

For Nef detection, the following mouse monoclonal antibodies were used: 3D12 (ABCAM ab42355) against amino acids ^35^RDLEKHGAITSSNTAA^50^ of HIV-1 HXB2; EH1 (AIDS reagents #ARP-3689) against ^194^MARELHPEYYKDC^206^ of HIV-1 B subtype consensus; SN20 (gift from Dr. Bernhard Maier, Indiana University) against the SH3 binding domain of Nef (FPVTPQ); and 6JR (ABCAM ab42358) mapped to ^195^ARELHPEYYKD^205^ of the HIV-1 B subtype consensus. P-Tau217 was detected using the rabbit polyclonal antibody to Tau phosphorylated on Threonine 217 (Thermofisher (Waltham, MA), cat. #44744); ABCA1 – using mouse monoclonal anti-ABCA1 antibody H10 (Abcam (Cambridge, MA), cat. #ab18180); and flotillin 1 – using goat anti-flotillin 1 polyclonal antibody (Novus Biologicals (Littleton, Co), cat. #NB1001043).

### Statistics

Statistical analyses including Spearman correlations, Wilcoxon rank-sum tests, Kruskal-Wallis test with DSCF multiple comparisons option, and multivariate linear regression were conducted in SAS v9.4. Data visualization was conducted in GraphPad PRISM v.9. For multivariate analysis, Box Cox transformation of variables was performed in SAS v9.4 prior to regression. Figure captions describe the statistical test used.

## Results

The role of amyloid proteins in pathogenesis of HAND has been previously suggested [23–25]. Our earlier study of a small number of postmortem brain samples from HAND patients found a trend towards increased abundance of APP and Tau relative to samples from uninfected individuals [11]. To substantiate this finding, we now analyzed postmortem brain samples from 22 HIV-infected individuals diagnosed with HAND, 11 HIV-infected individuals without HAND diagnosis, and 12 uninfected controls without diagnosed neurological disease (Table S1) for Tau protein phosphorylated on Threonine 217 (p-Tau217). This Tau isoform has been shown to strongly correlate with Aβ deposition [26] and was proposed as a marker for Alzheimer’s disease [20, 27, 28]. Results presented in Fig. 1A demonstrate that the abundance of p-Tau217, measured by quantitative automated capillary Western blot, was significantly increased in samples from HIV-infected individuals with HAND diagnosis, relative to HIV-infected individuals without HAND diagnosis and especially relative to uninfected controls (supporting Western blot evidence is presented in supplementary Fig. S1). Significant difference in the abundance of this peptide was also observed between samples from HIV-infected individuals without HAND diagnosis and uninfected controls (Fig. 1A). Given that p-Tau217 is an early marker of neurodegeneration and Alzheimer’s [20, 27, 28], these results are consistent with the role of phosphorylated Tau in HAND pathogenesis and suggest that increase in this factor may precede HAND diagnosis.

**Figure 1.**
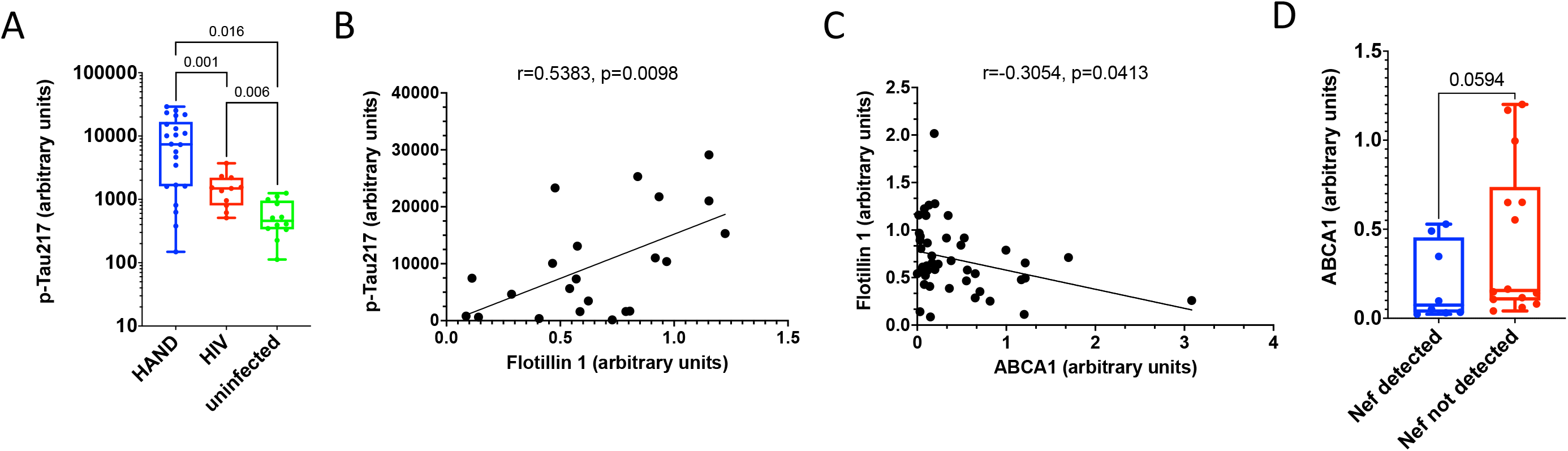
Analysis of pathogenic correlates of HAND. A – Analysis of p-Tau217 in brain samples from uninfected individuals and HIV-infected individuals with and without HA for p-Tau217 adjusted to total protein were obtained using ProteinSimple Compass software and are presented as arbitrary units. Group comparisons were made by Kruskal-Wallis test with post hoc Dwass, Steel, Critchlow-Fligner multiple comparison analysis adjustment for FWER (DSCF option in SAS NPAR1WAY procedure). B – Simple linear regression analysis of p-Tau217 and Flotillin 1 in HAND samples (data adjusted to total protein obtained using Compass software). C - Simple linear regression analysis of ABCA1 and Flotillin 1 in all samples (protein abundance adjusted to total protein was obtained using Compass software). D – Analysis of ABCA1 in HIV-infected (HAND-positive and HAND-negative samples) vs uninfected samples. Data points representing ABCA1 abundance adjusted to total protein were obtained using Compass and are presented as box and whiskers plot with p value calculated by Mann Whitney nonparametric two-tailed t test.

Amyloidogenic proteins and amyloid peptides are associated with lipid rafts [29–33], and our previous study suggested that increased abundance of lipid rafts caused by Nef-containing extracellular vesicles (exNef) may be the reason for the upregulation of APP and Tau [11]. Unfortunately, immunohistochemical analysis of lipid rafts in fresh-frozen tissue blocks was technically challenging. We evaluated the abundance of lipid rafts in the brain tissue samples by measuring the lipid raft marker flotillin 1 (by quantitative Western blot). Results of these analyses are presented in Fig. S2. Although there was no difference in the abundance of flotillin 1 between brain samples from HIV-infected HAND-positive or HAND-negative individuals vs uninfected individuals (Fig. S3), the abundance of p-Tau217 in HAND brain samples significantly correlated with the abundance of flotillin 1 (Fig. 1B). This result is consistent with the proposed relationship between lipid rafts and amyloid peptides [11, 32]. Of note, no such correlation was observed in samples from uninfected brains, or brains from HIV-infected individuals without HAND diagnosis (Fig. S4), suggesting that the relationship between rafts and p-Tau217 formation is changed in HAND.

Results above suggest that abundance p-Tau217 in HAND brains may be associated with the abundance of flotillin 1 and, by extension, of lipid rafts. Previous studies suggested that downmodulation of ABCA1 in macrophages infected with HIV-1 or treated with exNef regulates the abundance of lipid rafts [10–12, 34]. Our analysis showed a negative correlation between the quantity of ABCA1 in all samples and the quantity of flotillin 1 (Fig. 1C). Comparison of ABCA1 abundance between all samples from HIV-infected individuals to samples from uninfected controls showed a trend towards reduced ABCA1 in the brains from infected individuals, although the statistical significance was not achieved (Fig. 1D, supporting Western blot evidence is in Fig. S5).

A known problem with detection of Nef in biological samples is that Nef proteins from different HIV-1 strains have vastly varying ability to interact with different antibodies, significantly affecting accuracy of comparison of the abundance of Nef between infected participants. In addition, low specificity of most anti-Nef antibodies makes detection of low Nef concentrations in clinical samples challenging. To get around these limitations, we detected Nef in the brain samples by Western blot using four different antibodies raised against conservative epitopes in different Nef regions: 3D12 mapped to the N-terminus region of HXB2 Nef, EH1 mapped to the C-terminus region, SN20 mapped to the SH3-binding domain of Nef, and 6JR mapped to the C-terminus region (see Materials and Methods). Analysis with 3D12, SN20, and EH1 antibodies was performed with all samples, except HIV 40, HIV 42, HIV 43, HAND 41, HAND 44, HAND 45, and showed some cross-reactivity with cellular proteins in uninfected samples (Fig. S6). All samples were analyzed using the 6JR antibody, which provided the cleanest results, with least cross-reactivity with cellular proteins and negative results with all uninfected samples (Fig. 2), and was used for the final interpretation of the Nef status. Table 1 summarizes results of this analysis. Eight of 22 HAND samples were Nef-positive, whereas only one sample from HIV-positive individual without HAND diagnosis (HIV 19) was positive.

**Figure 2.**
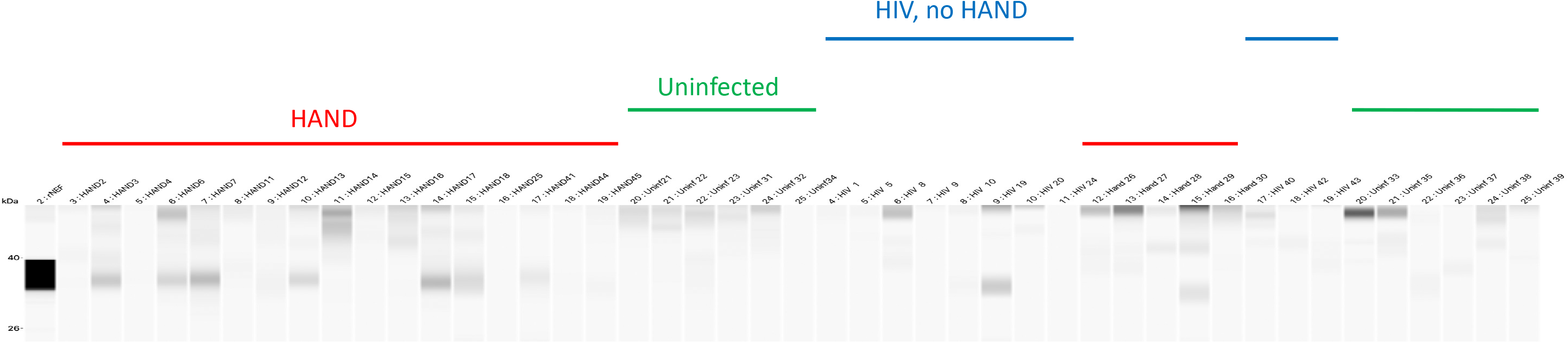
Analysis of Nef in brain samples. Brain lysates were assayed by Western immunoblot using the 6JR mouse monoclonal anti-Nef antibody from Abcam. Samples were run on ProteinSimple Jess instrument and analyzed by Compass software. Color-coded lines denote group assignment of the samples.

**Table 1.**
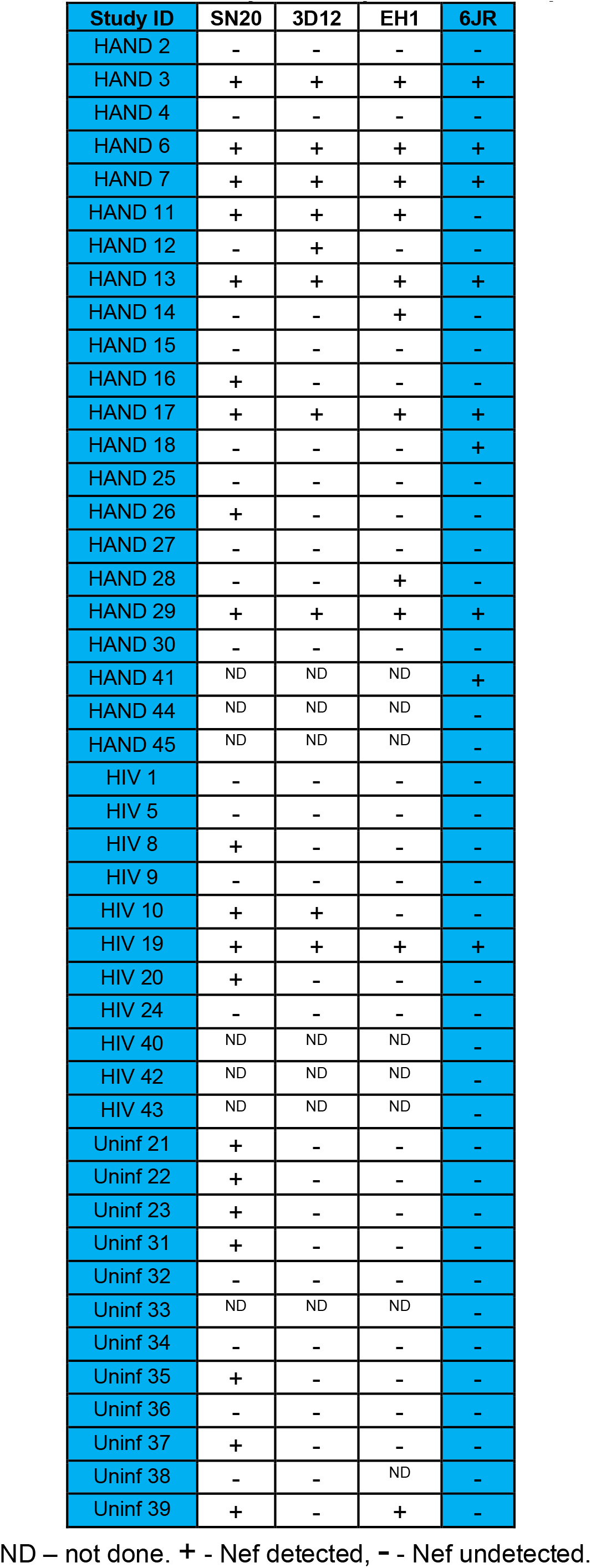
Results of Nef analysis in experimental samples.

We further separated HAND samples into two groups, Nef-positive and Nef-negative, and compared these groups for expression of ABCA1, flotillin 1, p-Tau217, and HAND severity according to the clinical score (Fig. 3). Samples positive for Nef had lower abundance of ABCA1 (Fig. 3A), higher abundance of flotillin (Fig. 3B), and higher abundance of p-Tau217 (Fig. 3C) relative to HAND samples with undetectable Nef. To determine whether the observed differences were associated with the disease status, we compared the clinical HAND scores between subjects providing the Nef-positive and Nef-negative brain samples. Clinical neurological status was assessed using three existing dementia rating scales, American Academy of Neurology (AAN), Memorial Sloan Kettering (MSK), and Frascati; the scales were not universally applied to all individuals and in some cases disagreed (Table S1). Although a good concordance between these scales has been reported, Frascati provided a more graded evaluation, especially for mild cases [35]. We therefore used the Frascati score, were available, to grade the clinical status, and the AAN score in the other cases (Table 2). The HAND clinical score was significantly higher in individuals providing Nef-positive samples relative to individuals providing samples with undetectable Nef (Fig. 3D), indicating that presence of Nef was associated with a more severe disease. Multivariate regression analysis demonstrated that the effect of detectable Nef on all continuous outcome variables (ABCA1, flotillin 1, p-Tau217) considered jointly was significant in all HIV-infected individuals, in both the unadjusted model and the adjusted model controlling for age, HIV duration, and viral load (Table S2). The difference in results between analysis of all samples from HIV-positive individuals and those from HAND only may be due to the smaller sample size in HAND only set.

**Figure 3.**
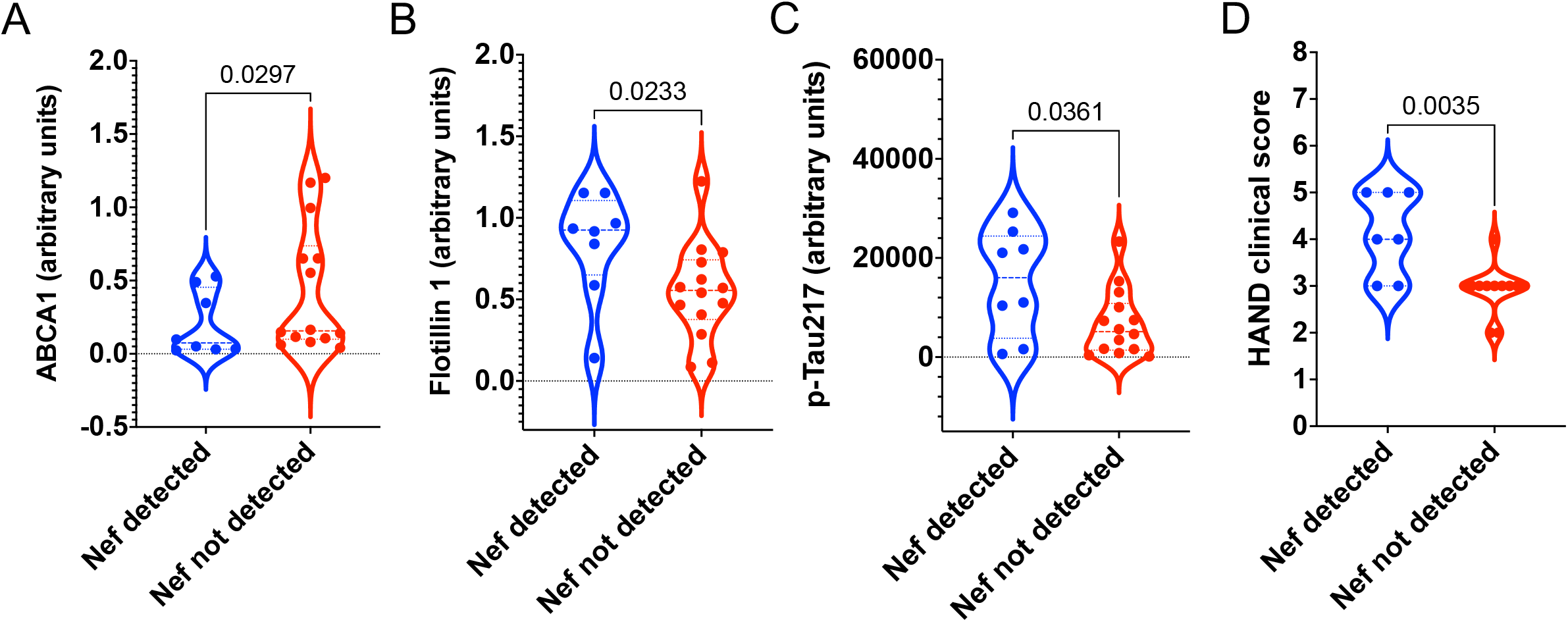
Comparative analysis of Nef-positive and Nef-negative brain samples from HAND-diagnosed individuals. A, B, C **-**Data points for ABCA1 (A), Flotillin 1 (B), and p-Tau217 (C) adjusted to total protein levels were obtained using ProteinSimple Compass software and are presented as arbitrary units. Results are presented as violin plots, showing p values calculated for unpaired parametric two-tailed t test. D – HAND clinical scores are presented for individuals from whom Nef-positive and Nef-negative brain samples were obtained as violin plots, showing p values calculated for unpaired non-parametric Mann-Whitney two-tailed t test.

**Table 2.** HAND severity evaluation in donors of post-mortem brain samples^A^.

| **ID** | **Frascati score** | **AAN score** | **HAND score in the study** |
| --- | --- | --- | --- |
| HAND 2 |  | 3 | 3 |
| HAND 3 |  | 5 | 5 |
| HAND 4 |  | 4 | 4 |
| HAND 6 |  | 4 | 4 |
| HAND 7 |  | 5 | 5 |
| HAND 11 |  | 3 | 3 |
| HAND 12 |  | 2 | 2 |
| HAND 13 |  | 5 | 5 |
| HAND 14 |  | 3 | 3 |
| HAND 15 |  | 3 | 3 |
| HAND 16 | 3 |  | 3 |
| HAND 17 | 3 |  | 3 |
| HAND 18 | 2 |  | 2 |
| HAND 25 | 3 |  | 3 |
| HAND 26 |  | 3 | 3 |
| HAND 27 |  | 3 | 3 |
| HAND 28 |  | 3 | 3 |
| HAND 29 |  | 3 | 3 |
| HAND 30 |  | 3 | 3 |
| HAND 41 | 5 | 4 | 5 |
| HAND 44 | 5 | 4 | 5 |
| HAND 45 | 3 | 5 | 3 |

<sup>A</sup>HAND score used in the study was based on Frascati score where available, and AAN score in other cases.

Taken together, these results suggest that Nef, via modification of cholesterol metabolism and lipid rafts may be one of the key causative factors promoting amyloid formation and disease progression in HAND.

## Discussion

The role of amyloid proteins in HAND pathogenesis remains an open question. In this study, we demonstrated a significant increase in the brains of HIV-infected individuals of the abundance of p-Tau217, which is also involved in pathogenesis of Alzheimer’s disease [36]. This finding is consistent with our previous report where *in vitro* analysis suggested increased abundance of amyloid precursor proteins and Tau in exNef-treated neuronal cells [11]. Surprisingly, this was true for brain samples from both HAND-diagnosed and HAND-free HIV-infected individuals. One explanation is that HIV-infected individuals without HAND diagnosis were at an early stage of disease when clinical manifestations could not yet be detected. This explanation is consistent with a demonstrated utility of p-Tau217 as an early marker of Alzheimer’s, which allows detection of the disease at the stage preceding neurodegeneration and clinical symptoms [37].

In these studies, we measured total flotillin 1 to evaluate the abundance of the lipid rafts. Flotillins 1 and 2 are integral membrane proteins that assist in raft assembly [38–40]. While flotillin 1 can be recruited to lipid rafts under certain stimulatory conditions [41], suggesting that the number of flotillin molecules per lipid raft may vary, the level of flotillin expression, and thus the per cell abundance of flotillin, appears to regulate the abundance of lipid rafts [42]. Therefore, the abundance of total flotillin 1 provides only a crude estimate of the abundance of lipid rafts. This limitation is taken in the account in our interpretations.

Tau hyperphosphorylation is a characteristic feature of Alzheimer’s disease, and a number of studies established that phosphorylation of Tau in AD is closely linked to Aβ pathology [26, 43, 44]. Lipid rafts were proposed to be the sites where Tau phosphorylation takes place, and where phosphorylated Tau accumulates and interacts with Aβ during development of Alzheimer’s disease [33]. The observed correlation between abundancies of flotillin 1 and p-Tau217 may reflect this paradigm, suggesting that lipid rafts may be a target for potential therapeutic interventions in HAND, as had been proposed for other neurodegenerative diseases [45]. We recognize, however, that, as discussed above, abundance of flotillin 1 provides only a crude estimate of the abundance of lipid rafts, which may be the reason for low flotillin 1 in some brain samples from subjects diagnosed with HAND, and high flotillin 1 in some samples from HIV-infected individuals without the HAND diagnosis. But the most likely explanation for this variability is that a driving factor in HAND pathogenesis is not the absolute amount, but the change in the quality and quantity of the lipid rafts during disease progression. Our previous studies documented functional impairment of lipid rafts associated with the effect of Nef [10], which may occur without a major change in the raft abundance. We have previously reported that exNef modify the fatty acid content of the lipid rafts [12], although the functional consequences of these changes have not been investigated. Prospective studies will be needed to evaluate this idea.

Our analysis demonstrated that majority of samples where Nef could be detected came from individuals with diagnosed HAND. This result suggests an important role of Nef in HAND pathogenesis. Indeed, Nef-positive samples were characterized by reduced ABCA1, increased flotillin 1, increased p-Tau217, and increased clinical disease score. These results suggest that Nef may drive ABCA1 downmodulation in HAND initiating lipid raft modifications and subsequent pathological events.

A limitation of this study is a relatively small number of samples available for analysis. Since samples were not cryopreserved, we could not perform flow cytometry analysis of the lipid rafts, or cell-specific characterizations. Another limitation is lack of a sensitive quantitative assay for Nef. Despite these limitations, our study supports the pathogenic role of Tau in HAND pathogenesis, and suggests the sequence of events that lead to HAND: Nef EVs downmodulate ABCA1 changing the properties of the lipid rafts, thus increasing formation of amyloid plaques and Tau phosphorylation and fibrillation. Future studies in animal models will establish the temporality of this ordering and will determine whether therapeutic treatments breaking the pathogenic course described above can prevent development of HAND.

## Author Declarations

### Ethics approval and consent to participate

Not applicable. Anonymized samples were received from repositories.

### Consent for publication

All authors received the final version of the manuscript and agreed to publish it.

### Availability of data and materials

All the data are presented in the manuscript as the main or supplementary material. The sources of materials are provided.

### Authors’ contributions

TP performed the experiments and contributed to data analysis and interpretation; AW performed statistical analysis; DS contributed to research design, results’ interpretation, and manuscript preparation; MIB designed the experiments, contributed to results’ interpretation, wrote the manuscript.

## Acknowledgements

We are thankful to Drs. Matthias Clauss and Bernhard Maier (Indiana University and Purdue University Indianapolis) for anti-Nef antibody SN20. The following reagent was obtained through the NIH HIV Reagent Program, Division of AIDS, NIAID, NIH: Anti-Human Immunodeficiency Virus 1 (HIV-1) Nef Monoclonal (EH1), ARP-3689, contributed by Dr. James Hoxie. We are thankful to NIH NeuroBioBank for providing brain samples from uninfected individuals. The brain samples from HIV-infected individuals with and without HAND diagnosis were provided by The National NeuroAIDS Tissue Consortium (NNTC) supported by shared resources from NIH funding through the NIMH and NINDS by the following grants: Manhattan HIV Brain Bank (U24MH100931), Texas NeuroAIDS Research Center (U24MH100930) National Neurological AIDS Bank (U24MH100929), California NeuroAIDS Tissue Network (U24MH100928), Data Coordinating Center (U24MH100925).

## Compliance with Ethical Standards

### Research involving Human Participants and/or Animals

Human samples analyzed in this study were anonymized and were provided by NIH-maintained repositories.

### Informed consent

Not applicable.

