## Supplemental Table 1 for "Abundance of Nef and p-Tau217 in brains of individuals diagnosed with HIV-associated neurocognitive disorders correlate with disease severance"

**Table S1. Clinical data for the individuals from whom brain samples were obtained.**

| Original ID[A] | Study ID[B] | Age[A] | Gender[A] | Frascati[A] | AAN[A] | MSK[A] | VL[A] | logVL[A] | HIV duration[A] |
|---|---|---|---|---|---|---|---|---|---|
| 1071 | HAND 2 | 39 | M | | 3 | 1 | 8584 | 3.93 | 12 |
| 4041 | HAND 3 | 34 | F | | 5 | 2 | 165682 | 5.22 | 5 |
| 4028 | HAND 4 | 46 | F | | 4 | 2 | 750000 | 5.88 | 6 |
| 6011 | HAND 6 | 49 | M | | 4 | 2 | 5000 | 3.7 | 10 |
| 6046 | HAND 7 | 50 | M | | 5 | 2 | 400 | 2.6 | 15 |
| 7100118382 | HAND 11 | 37 | M | | 3 | 2 | 178 | 2.25 | 19 |
| 1021 | HAND 12 | 41 | F | | 2 | 1 | 400 | 2.6 | 5 |
| 2066 | HAND 13 | 35 | M | | 5 | 2 | 250000 | 5.4 | 13 |
| 1159 | HAND 14 | 44 | M | | 3 | 1 | 69931 | 4.84 | 13 |
| 2074 | HAND 15 | 36 | M | | 3 | 1 | 6952 | 3.84 | 3 |
| CE135 | HAND 16 | 39 | M | 3 | | 1 | 198779 | 5.3 | 13 |
| CE116 | HAND 17 | 46 | F | 3 | | | 342386 | 5.53 | 13 |
| CC101 | HAND 18 | 44 | M | 2 | | | 39530 | 4.6 | 19 |
| CC202 | HAND 25 | 36 | M | 3 | | 1 | 750000 | 5.88 | |
| 7100056883 | HAND 26 | 35 | M | | 3 | 1 | 18985 | 4.28 | 6 |
| 7100686683 | HAND 27 | 31 | M | | 3 | 2 | 483758 | 5.68 | 5 |
| 7200077165 | HAND 28 | 45 | M | | 3 | 1 | 316227 | 5.5 | 8 |
| 7200116866 | HAND 29 | 36 | M | | 3 | 2 | 433715 | 5.64 | 5 |
| 7200017177 | HAND 30 | 41 | M | | 3 | 2 | 501 | 2.7 | 16 |
| 010123 | HAND 41 | 44 | F | 5 | 4 | 2 | 129083 | 5.11 | 13 |
| 030013 | HAND 44 | 30 | M | 5 | 4 | 2 | 104300 | 5.02 | 8 |
| 030025 | HAND 45 | 46 | M | 3 | 5 | 2 | 750000 | 5.88 | 3 |
| 6052 | HIV 1 | 39 | M | | 0 | | 75000 | 4.88 | 14 |
| 4109 | HIV 5 | 32 | M | | 0 | | 400 | 2.6 | 4 |
| CE122 | HIV 8 | 39 | M | 0 | | | 229168 | 5.36 | 20 |
| CB194 | HIV 9 | 37 | M | 0 | | | 9542 | 3.98 | 9 |
| 7200207472 | HIV 10 | 35 | M | | 0 | | 135643 | 5.13 | 5 |
| CA236 | HIV 19 | 34 | M | 0 | | | 552000 | 5.74 | 8 |
| CC163 | HIV 20 | 45 | M | 0 | | | 16700 | 4.22 | 21 |
| CE119 | HIV 24 | 41 | F | 0 | | | 83061 | 4.92 | 11 |
| 010104 | HIV 40 | 50 | M | 0 | 3 | | 26621 | 4.43 | 14 |
| 010251 | HIV 42 | 40 | F | 0 | 4 | | 77 | 1.89 | 17 |
| 030006 | HIV 43 | 37 | M | 0 | | | 287947 | 5.46 | 4 |
| GW5 | uninf 21 | 55 | F | | | | | | |
| GW22 | uninf 22 | | F | | | | | | |
| GW367 | uninf 23 | 84 | M | | | | | | |
| HU13350 | uninf 31 | 45 | M | | | | | | |
| HU13134 | uninf 32 | 53 | M | | | | | | |
| HU13477 | uninf 33 | 58 | M | | | | | | |

| | | | |
|---|---|---|---|
| HU13345 | uninf 34 | 62 | M |
| HU13440 | uninf 35 | 68 | M |
| HU13323 | uninf 36 | 24 | F |
| HU13344 | uninf 37 | 33 | F |
| HU13243 | uninf 38 | 53 | F |
| HU13114 | uninf 39 | 54 | F |

[A] Original sample IDs. Samples from HIV-infected individuals with or without HAND diagnosis were provided by NNTC, samples from uninfected individuals came from GWU (GW) or NBB (HU). All clinical data were supplied by the providers. VL – viral load (RNA copies/ml), HIV duration – known length of HIV infection (years).
[B] IDs used in the study. HAND or HIV (no HAND) designation was made based on NNTC grouping.
