## Supplemental Table 2 for "Abundance of Nef and p-Tau217 in brains of individuals diagnosed with HIV-associated neurocognitive disorders correlate with disease severance"

| **Table S2. Multivariate linear regression results for Nef (undetectable vs. detectable) predicting outcomes** | | | | | | | | | | |
|---|---|---|---|---|---|---|---|---|---|---|
| **Outcome Variable[A]** | **Estimate** | **95% LCL** | **95% UCL** | **p-Value** | **MANOVA p-Value[B]** | **Adjusted Estimate[C]** | **Adjusted 95% LCL** | **Adjusted 95% UCL** | **Adjusted p-Value** | **Adjusted MANOVA p-Value** |
| *Participants with HAND (n=22)* | | | | | | | | | | |
| ABCA1[D] | -0.22 | -0.49 | 0.05 | 0.105 | - | -0.15 | -0.46 | 0.17 | 0.331 | - |
| Flotillin-1 | 0.26 | -0.02 | 0.54 | 0.070 | - | 0.19 | -0.16 | 0.53 | 0.271 | - |
| p-Tau217[E] | 0.48 | -0.08 | 1.04 | 0.089 | - | 0.33 | -0.24 | 1.14 | 0.183 | - |
| Overall model | - | - | - | - | 0.051 | - | - | - | - | 0.304 |
| *All participants with HIV (n=33)* | | | | | | | | | | |
| ABCA1[D] | -0.14 | -0.39 | 0.10 | 0.244 | - | -0.10 | -0.37 | 0.16 | 0.423 | - |
| Flotillin-1 | 0.10 | -0.21 | 0.40 | 0.511 | - | 0.08 | -0.25 | 0.41 | 0.634 | - |
| p-Tau217[E] | **0.62** | **0.20** | **1.04** | **0.005** | - | **0.65** | **0.20** | **1.09** | **0.006** | - |
| Overall model | - | - | - | - | **0.014** | - | - | - | - | **0.030** |
| [A]Modeling effect of the predictor Nef = detectable (undetectable as the reference category) | | | | | | | | | | |
| [B]MANOVA p-value by Pillai's trace exact F test for the overall effect of Nef | | | | | | | | | | |
| [C]Controlling for age, HIV duration (in years), and $log_{10}$ viral load | | | | | | | | | | |
| [D]Modeling square root of ABCA1 values | | | | | | | | | | |
| [E]Modeling $log_{10}$ p-Tau217 values | | | | | | | | | | |
| Note: significant effects at the alpha=0.05 level are bolded | | | | | | | | | | |
| LCL - lower confidence limit; UCL - upper confidence limit; MANOVA - multivariate analysis of variance | | | | | | | | | | |
