## Supplemental Figure 1 for "Abundance of Nef and p-Tau217 in brains of individuals diagnosed with HIV-associated neurocognitive disorders correlate with disease severance"

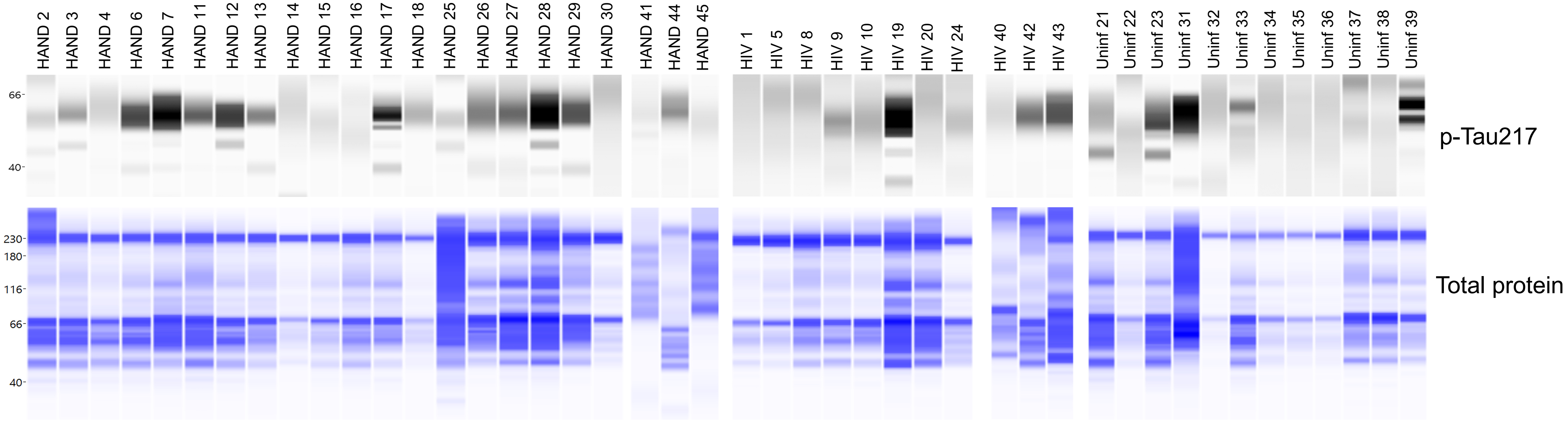


**Figure S1. Western blots for p-Tau217 and total proteins.** Molecular weight markers are shown on the left.
