## Supplemental Figure 2 for "Abundance of Nef and p-Tau217 in brains of individuals diagnosed with HIV-associated neurocognitive disorders correlate with disease severance"

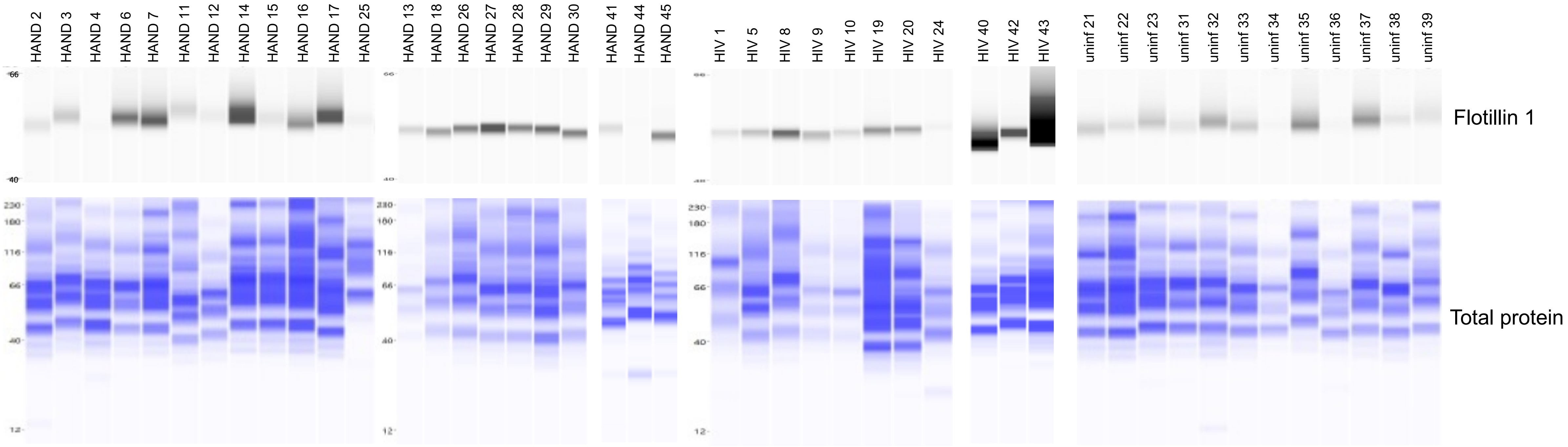


**Figure S2. Western blots for Flotillin 1 and total proteins.** Molecular weight markers are shown on the left.
