## Supplemental Figure 3 for "Abundance of Nef and p-Tau217 in brains of individuals diagnosed with HIV-associated neurocognitive disorders correlate with disease severance"

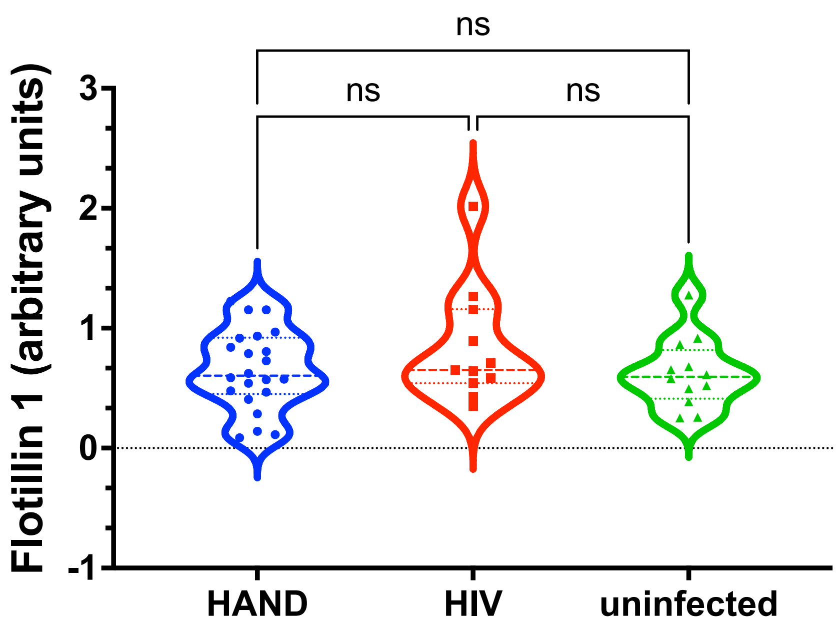


**Figure S3. Comparison of Flotillin 1 abundance between brain samples from HIV-infected individuals with (HAND) and without (HIV) HAND diagnosis and uninfected individuals.** Data points for Flotillin 1 adjusted to total protein levels were obtained using ProteinSimple Compass software and are presented as arbitrary units. Results are presented as violin plots, p values were calculated using Kruskal-Wallis test.
