## Supplementary figures and images for "Abundance of Nef and p-Tau217 in brains of individuals diagnosed with HIV-associated neurocognitive disorders correlate with disease severance"

### Supplemental Figure 4

$r^2$=0.008, p=0.794

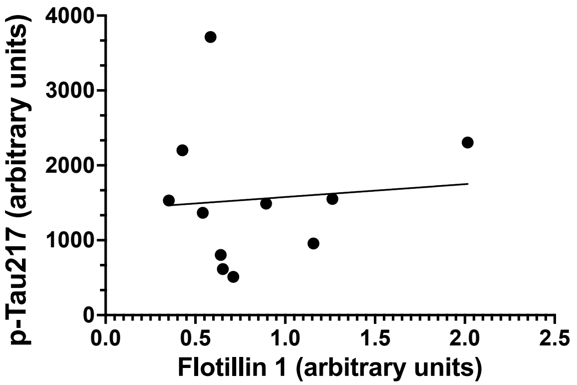


HIV-infected, no HAND

$r^2$=0.005, p=0.821

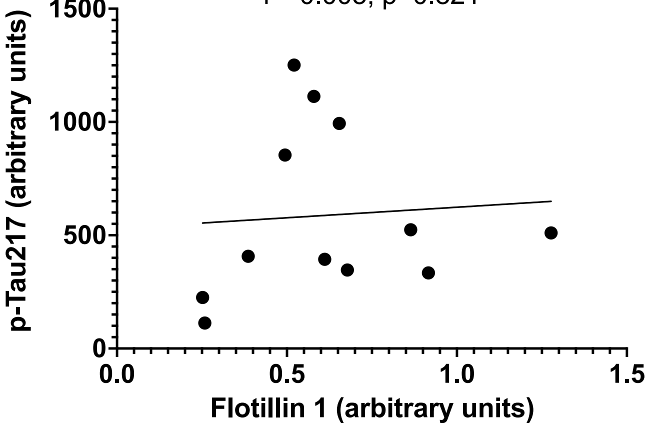


uninfected
