## Supplemental Figure 6 for "Abundance of Nef and p-Tau217 in brains of individuals diagnosed with HIV-associated neurocognitive disorders correlate with disease severance"

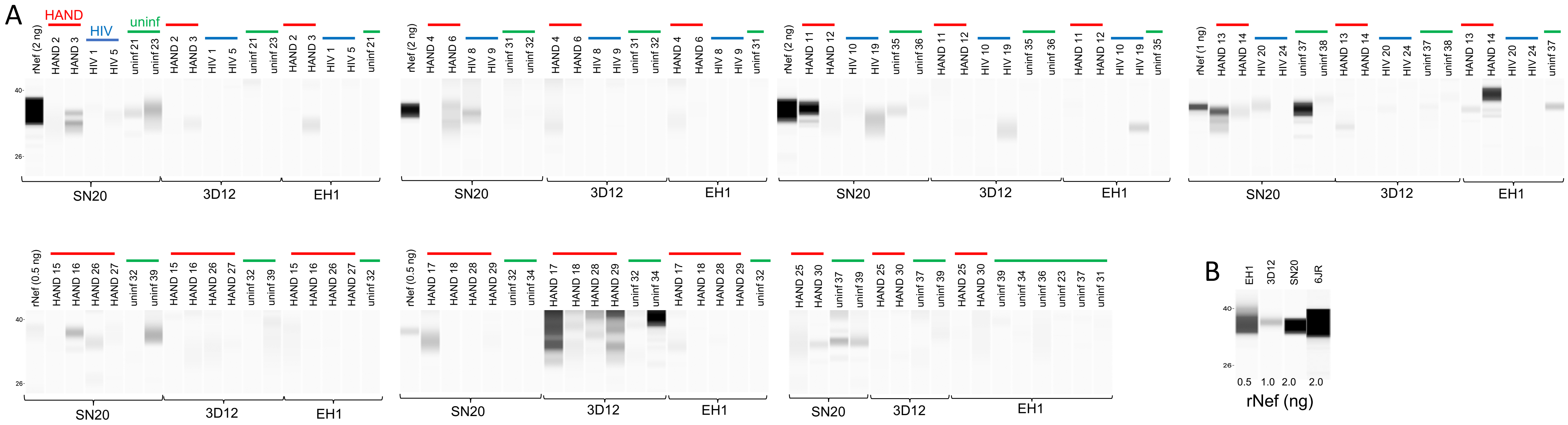


**Figure S6. Analysis of Nef in brain samples.** Brain lysates were analyzed on ProteinSimple Jess instrument using the Compass software. A – Sample IDs are shown on top of the gels, and antibodies used for detection – below the gels. B – Detection of recombinant Nef from HIV-1$_{SF2}$ by antibodies used in this study. Color coded lines show the sample group.
